# A Light sheet fluorescence microscopy and machine learning-based approach to investigate drug and biomarker distribution in whole organs and tumors

**DOI:** 10.1101/2023.09.16.558068

**Authors:** Niyanta Kumar, Petr Hrobař, Martin Vagenknecht, Jindrich Soukup, Nadia Patterson, Peter Bloomingdale, Tomoko Freshwater, Sophia Bardehle, Roman Peter, Ruban Mangadu, Cinthia V. Pastuskovas, Chiswili Y. Chabu, Mark T. Cancilla

## Abstract

Tissue clearing and Light sheet fluorescence microscopy (LSFM) provide spatial information at a subcellular resolution in intact organs and tumors which is a significant advance over tools that limit imaging to a few representative tissue sections. The spatial distribution of drugs, targets, and biomarkers can help inform relationships between exposure at the site of action, efficacy, and safety during drug discovery. We demonstrate the use of LSFM to investigate distribution of an oncolytic virus (OV) and vasculature in xenograft tumors, as well as brain Aβ pathology in an Alzheimer’s disease (AD) mouse model. Machine learning-based image analysis tools developed to segment vasculature in tumors showed that random forest and deep learning methods provided superior segmentation accuracy vs intensity-based thresholding. Sub-cellular resolution enabled detection of punctate and diffuse intracellular OV distribution profiles. LSFM investigation in the brain in a TgCRND8 AD mouse model at 6.5 months of age enabled evaluation of Aβ plaque density in different brain regions. The utility of LSFM data to support quantitative systems pharmacology (QSP) and physiology-based pharmacokinetics (PBPK) modeling to inform drug development are also discussed. In summary, we showcase how LSFM can expand our understanding of macromolecular drug and biomarker distribution to advance drug discovery and development.

## Introduction

Assessing the spatial distribution of macromolecular drugs, targets, and biomarkers can provide critical information to advance drug discovery and development. Most high-resolution *ex vivo* optical microscopy methods can only image up to a limited depth within tissues of interest (typically ∼50-100 µm for confocal microscopy and up to ∼1 mm for two photon microscopy).

This is due to a proportional increase in light scattering and the diffusion barrier for molecular probes with increasing depth within tissues, which is primarily attributed to the tightly packed lipid bilayer of cell membranes. As a result of throughput constraints, analyses are typically restricted to a few representative thin tissue sections that do not fully capture the heterogeneity in whole organs and tumors, subsequently limiting the robustness and accuracy of the quantitative information obtained (**Fig. 1**). In this work, we demonstrate the application of a tissue clearing, immunolabeling, and light sheet fluorescence microscopy (LSFM) workflow ^1, 2^ for rapidly acquiring macromolecular drug, anatomical, and pathological marker tissue distribution information in intact organs and tumors without compromising on spatial resolution (**Fig. 2**).

**Figure 1.**
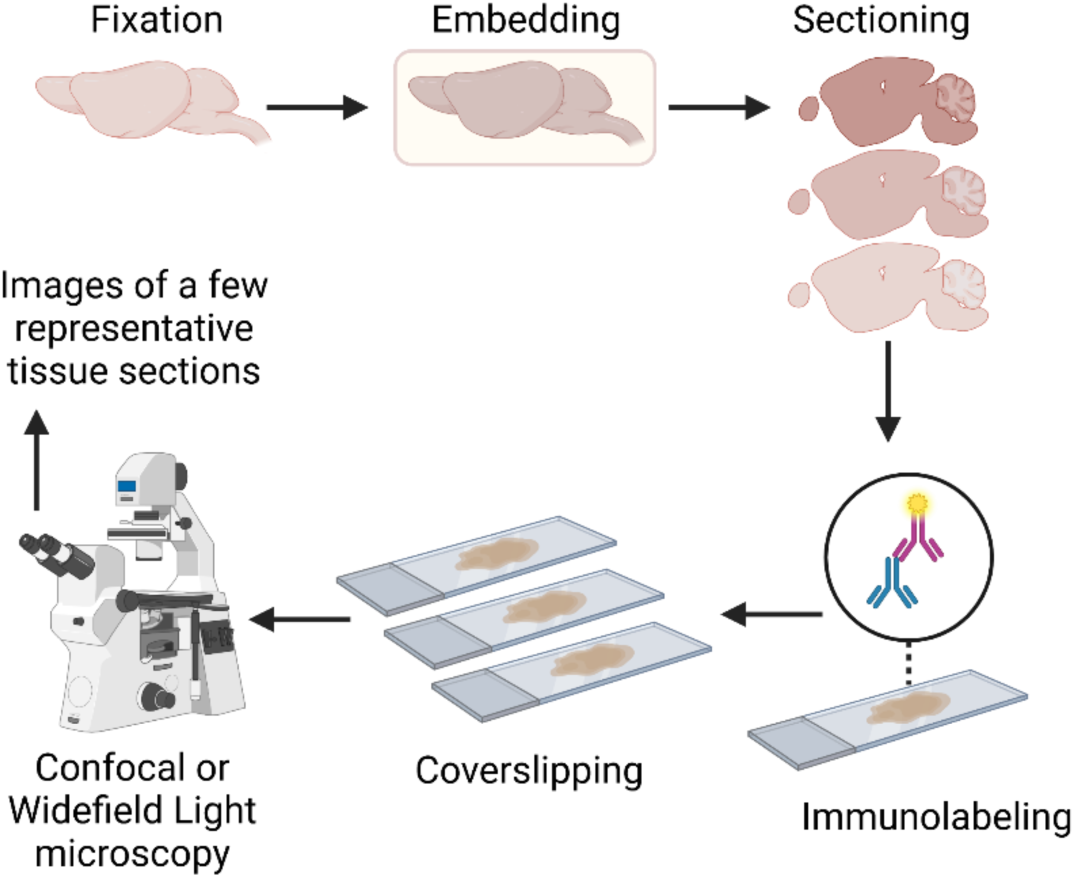
Schematic of a typical histology workflow requiring sectioning tissue samples into thin (typically 10-50 μm thick) sections, immunolabeling, then imaging with only a few representative sections. Created with BioRender.com.

**Figure 2.**
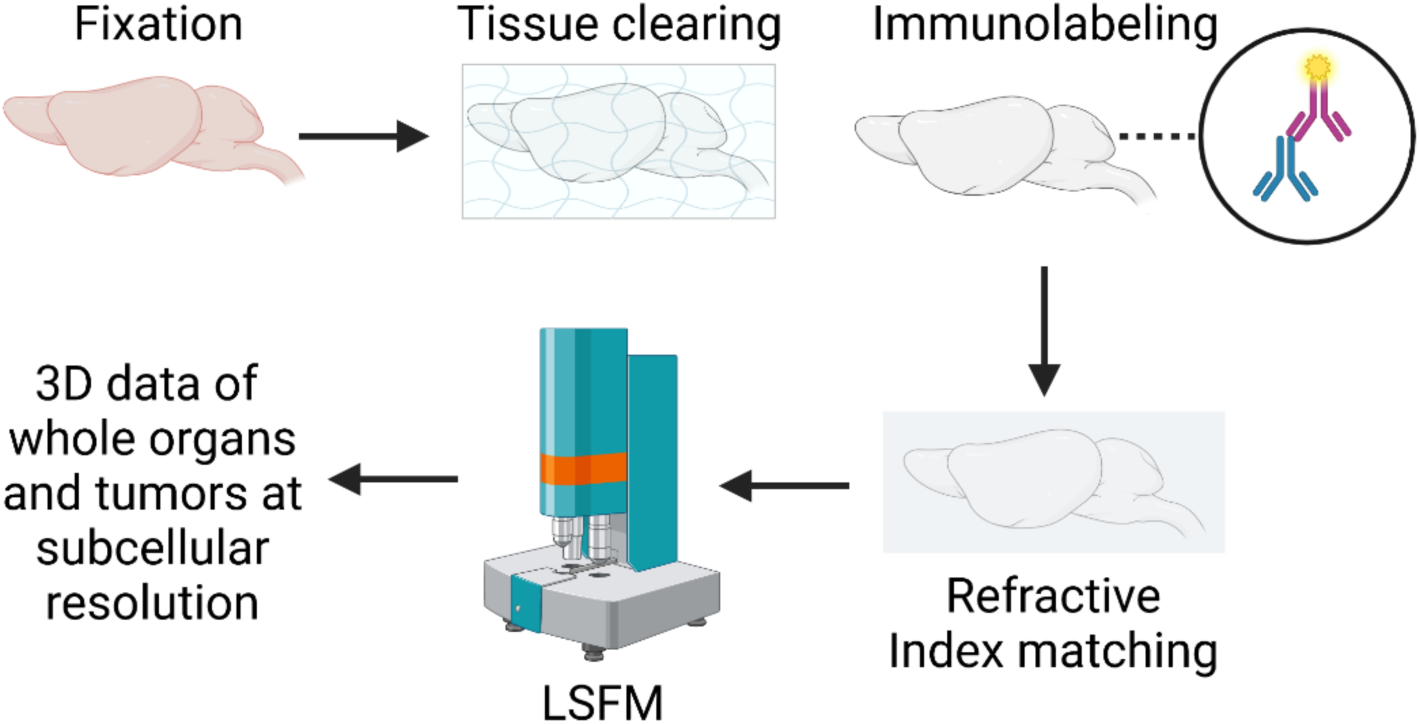
Tissue clearing and LSFM workflow: Tissue preservation using aldehyde and epoxy-based fixation. Tissue clearing with gentle delipidation while other macromolecules like proteins, DNA, and RNA are preserved. Performed using a surfactant-containing buffer and in an instrument supporting rotational electrophoresis for greater speed and uniformity versus passive diffusion. Immunolabeling via rotational electrophoresis. Refractive Index matching of cleared tissue to immersion liquid, rendering tissue optically transparent followed by light sheet fluorescence microscopy. Created with BioRender.com.

We utilized an aqueous tissue clearing method developed by Chung et al.,^3^ where a clear hydrogel-based framework is used to non-destructively replace lipids in fixed tissue while maintaining the location and identity of protein and nucleotides. Prior to tissue clearing, SHIELD (Stabilization under Harsh conditions via Intramolecular Epoxide Linkages to prevent Degradation) reagents were used to preserve native protein antigenicity, nucleotide transcripts, and tissue architecture ^4^. The process also applied SWITCH (System-Wide control of Interaction Time and kinetics of CHemicals) technology that allows synchronizing both the tissue preservation reaction and the labeling reactions to improve probe penetration depth and uniformity of staining ^5^. The SmartClear and SmartLabel instruments and reagents (from Life Canvas Technologies) were utilized. These instruments employ stochastic electrotransport, a novel electrokinetic method that uses a rotational electric field to selectively disperse highly electromobile molecules through a porous sample without displacing the low-electromobility molecules ^6^. This enables completely clearing preclinical rodent organs/tumors and staining them with nuclear dyes and fluorophore tagged antibodies, overcoming the long timelines of aqueous tissue clearing and labeling methods for intact organs and tumors and limitations of depending on passive diffusion alone. Volumetric imaging of intact cleared and labeled brains and tumors was performed using LSFM. LSFM entails scanning the cleared tissue sample with a laser light sheet of adjustable wave-length which excites fluorophores in the plane of illumination and results in 2D cross-sectional fluorescence images through the entire depth of the tissue sample producing a volumetric 3D dataset ^7^. The tissue clearing and LSFM workflow is summarized in **Fig. 2**.

The first use-case demonstrates the use of LSFM to characterize tumor vasculature and penetration of an oncolytic virus, coxsackievirus A21 (CVA21) ^8^ in xenograft tumors. Examining the distribution of biologics within tumors in the context of vasculature can help understand the relationship between drug and vascular distribution. Using the work by Dobosz et al. ^9^ as an example we share our own machine-learning based image analysis methods developed using Python to enable extraction of quantitative readouts such as tumor vascular volume fraction and biologic penetration into tissue. The modular nature of our code makes it easy to modify and re-purpose the application for future LSFM or similar 3D imaging data analysis.

The second use-case demonstrates the application of tissue clearing and LSFM to examine the distribution of beta amyloid pathology in the TgCRND8 mouse model of Alzheimer’s disease ^10^. We showcase an application of the Smart Analytics platform from Life Canvas Technologies that supports image segmentation to allow amyloid plaque detection, followed by quantification and registration with the Allen brain atlas for visualization of metrics in the context of discrete brain regions. Finally, we discuss critical considerations when onboarding a 3D histology platform in a biopharmaceutical industry setting and the potential utility of such data in building predictive *in silico* models of disease progression.

### Methods

## 1. Examining drug and vascular distribution in intact xenograft tumors using tissue clearing, immunolabeling, and LSFM

### 1.1. Animal model, biologic dosing, and tissue collection

All animal experimental protocols were approved by the Institutional Animal Care and Use Committee at Merck & Co., Inc., Rahway, NJ, USA and were carried out in accordance with the NIH Guide for the Care and Use of Laboratory Animals ^11^. Severe combined immune-deficient (SCID) mice were inoculated with SK-MEL-28 human myeloma cells and mice with tumor size of ∼ 100 mm^3^ were selected for dosing. Selected mice received 10^7 TCID50 (Median Tissue Culture Infectious Dose) of CVA21 in saline dosed intravenously. At 24 hours post dosing, mice were exsanguinated by transcardial perfusion with 20 mL ice-cold 1X phosphate buffered saline (PBS) with 10 U/ml heparin or until perfusate ran clear, followed by 20 mL ice-cold 4% paraformaldehyde (PFA) at 5 ml/min flow rate. Transcardial perfusion was performed as previously described by Gage et al ^12^. The 4% PFA solution was freshly prepared by diluting 32% Paraformaldehyde Solution (15714-S Electron Microscopy Sciences) with 1X PBS. Tumors were excised and placed in 4% PFA at 4°C for post fixation for 24 hours. Tumor samples were then transferred to 1X PBS and stored at 4°C. An n = 3 tumor samples from separate animals were cleared, labeled, imaged, and analyzed using the method described below.

### 1.2. Fixation, Tissue Clearing, Labeling, and Refractive Index matching

Tissue processing involved 4 main steps: SHIELD Fixation, Tissue Clearing, Labeling, and Refractive Index matching per the protocol provided by Life Canvas Technologies (LCT).

#### 1.2.1. SHIELD Fixation

SHIELD OFF solution containing 50% SHIELD-Epoxy Solution (Part # SH-ES, LCT), 25% SHIELD Buffer Solution (Part # SH-BS, LCT), and 25% deionized (DI) water was freshly prepared, mixed by vortexing after addition of each component to prevent precipitate formation, and placed on ice. Tumor samples were transferred to the freshly prepared SHIELD OFF Solution (20 ml per tumor) and incubated at 4°C with shaking for 3 days. Tumor samples were then transferred to and incubated in SHIELD-ON Buffer (Part # SH-ON, LCT), 20 ml per sample, for 24 hours, at 37°C with shaking, completing the SHIELD preservation procedure. Tumor samples were transferred to 1X PBS with 0.02% sodium azide and stored at 4°C until the next step.

#### 1.2.2. Tissue Clearing

Clearing of tumor samples was performed on the SmartClear II Pro device from LCT per vendor protocol. Tumor samples were first incubated in Clearing Buffer A (LCT) overnight at 37°C on a rocker. Next, samples were placed in mesh bags which were then inserted in the SmartClear II Pro sample holder. Tissue clearing was performed on gentle mode (42° C, 1500 mA) until samples were homogenously translucent (∼10-15 days depending on tumor size). Following tissue clearing, sodium dodecyl sulfate (SDS) from the samples was washed out with incubation in 1X PBS at room temperature overnight on a rocker before transferring to fresh 1X PBS and storage at 4°C.

#### 1.2.3. Labeling

A sequential labeling protocol was applied, and tumors were labeled using the SmartLabel device (LCT) based on vendor recommendations. Tumor samples were first incubated in Primary Sample Buffer from LCT (∼ 20 ml per wash) in a conical tube overnight at room temperature (RT) with shaking. Samples were switched to fresh Primary Sample Buffer for an additional 4-hour incubation at room temperature with shaking. Primary antibody cocktail was prepared in 1 ml fresh Primary Sample Buffer with the following components:

- Rabbit anti-CVA21 primary antibody (Viralytics, Batch #NBF0204.03) - polyclonal purified IgG fraction of antiserum raised in rabbits against a purified preparation of Coxsackievirus A21. 7 mg/ml in Dulbecco’s Phosphate Buffered Saline (pH 7.5). Final working concentration 7 µg/ml.
- Goat anti-mouse CD31 IgG primary antibody (Novus Biologicals, Catalog no. AF3628). Detects mouse CD31/PECAM-1 and serves as an endothelial cell marker for blood vessels. Final working concentration 7 µg/ml.
- Syto16 – a fluorescent DNA intercalating dye – was also added to the primary antibody cocktail to label cell nuclei (Thermo Fisher, Catalog no. S7578). Final working concentration 8 µM.

Primary antibody labeling was performed in the SmartLabel device from LCT for 22 hours. Current on the electrophoresis chamber was ∼ 135 mA at start and increased gradually during the labeling process. Following labeling, the SmartLabel device reservoirs were drained, and Primary Sample Buffer was replaced with fresh buffer. The electrophoresis on the SmartLabel device was run for 2 hours. After 2 hours, the sample cup from the labeling chamber, was drained, washed and filled with Sample Wash Buffer (LCT). The sample cup was placed back into the SmartLabel device and electrophoresis was turned on to wash the tumor samples for 1 hour. Sample washes were repeated three times. Next, secondary antibody labeling was performed on the tumor samples. A secondary antibody cocktail was prepared in 1 ml Sample Wash Buffer with the following components:

- AlexaFluor 594 labeled donkey anti-rabbit IgG secondary antibody (Thermo Fisher, Catalog no. A32754). Final working concentration: 4.4 µg/ml.
- Alexa Fluor 642 labeled donkey anti-goat IgG secondary antibody (Thermo Fisher, Catalog no. A-21447). Final working concentration: 5 µg/ml.

Secondary antibody labeling was then performed using the SmartLabel device (LCT). The sample cup from the labeling chamber was first drained and 3 ml fresh Sample Wash Buffer with the sample was added to it. The secondary antibody cocktail (1 ml) prepared in the previous step was added to the sample cup and mixed by pipetting. The sample cup was placed back in the labeling chamber and electrophoresis was run for 6 hours with sample rotation turned on. After labeling, the sample cup was removed from the labeling chamber, washed, and filled with fresh Sample Wash Buffer. Three sample washes for 1 hour each were performed following the same procedure as the washes after primary antibody labeling. Tumor samples were next washed in 0.5X PBS (50% 1X PBS and 50% distilled water) to wash out unbound probes. Following a 4-hour incubation in 0.5X PBS, tumor samples were transferred to 1X PBS and incubated for 3 hours more to complete washing of any remaining unbound probes. Samples were then incubated in 4% PFA in 1X PBS overnight at RT with shaking to fix immunolabeling. Finally, samples were washed with 1X PBS at RT with shaking for at least 6 hours, refreshing the solution at least 3 times to wash out any PFA. Tumor samples were then stored in 1X PBS at 4°C until refractive index matching.

#### 1.2.4. Refractive Index Matching

Tumor samples were incubated in fresh 50% EasyIndex (EI) solution (LCT) diluted in distilled water for 24 hours with shaking at 37°C. Samples were then transferred into 100% EI and incubated for 24 hours with shaking at 37°C.

### 1.3. Tumor LSFM

Refractive index matched tumor samples were mounted on the Selective Plane Illumination Microscope (SPIM) microscope (SmartSPIM, LCT). Mounted samples were allowed to shake in EI solution at 37°C for 2 hours and then the sample was allowed to float in the SPIM sample chamber overnight to allow full equilibration before image acquisition. Image acquisition was performed using the 3.6X objective (xy pixel size: 1.8 μm, z-step: 4 μm) using the 488, 561, and 642 nm lasers at laser powers of 40%, 70%, and 70% respectively.

### 1.4. Tumor LSFM data visualization

2D raw data TIFF files were transformed using the Imaris file converter (Bitplane) into a single .ims 3D data file. Voxel size was set to (1.8 µm x 1.8 µm x 4 µm) to match the SPIM image acquisition settings. 3D stitched data was then visualized in Imaris software from Bitplane (version 9.7.2).

### 1.5. Tumor LSFM data analysis

To obtain estimated biologic penetration in tumors in the context of vasculature we developed an image analysis pipeline broadly based on prior work by Dobosz et al. ^9^ In this paper, we share our modular Python-based image analysis approach. Our light sheet image analysis pipeline consisted of four main steps – blood vessel segmentation, tumor boundary segmentation, vascular distance map creation, and data aggregation, respectively (**Fig. 3**). In this section, we describe each step of the pipeline individually. The data analysis pipeline code can be found in our GitHub repository (https://github.com/Merck/3D_Tumor_Lightsheet_Analysis_Pipeline), which includes an interactive demo notebook (https://github.com/Merck/3D_Tumor_Lightsheet_Analysis_Pipeline/blob/main/demo_script.ipynb).

**Figure 3.**
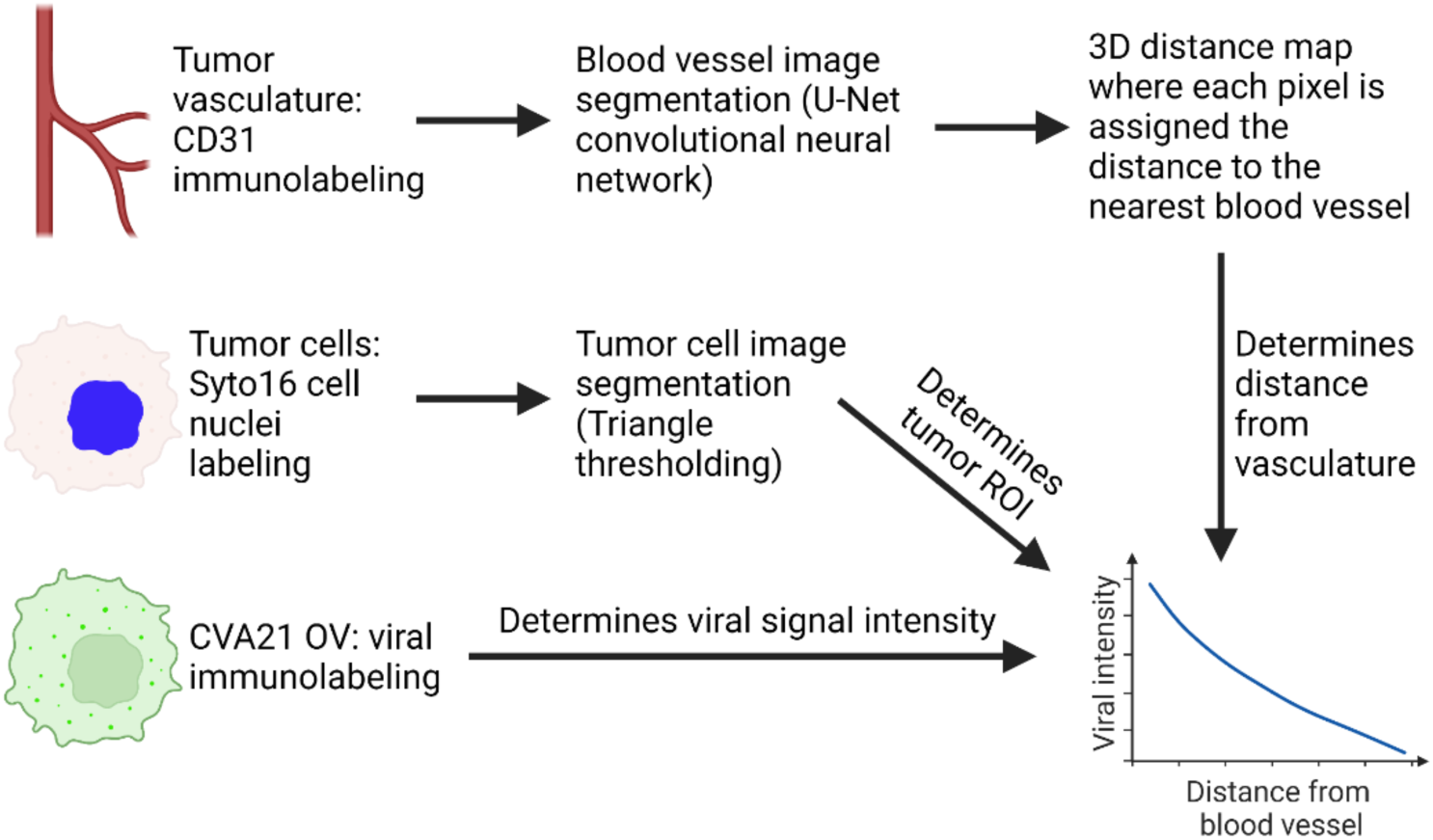
Schematic describing the tumor LSFM data analysis pipeline. Created with BioRender.com.

#### 1.5.1. Blood vessel segmentation

3D image segmentation of complex vasculature in large datasets is relatively uncharted territory. Filter-based methods such as intensity-based thresholding are a commonly used approach in image segmentation ^13^. We tested manual and automated (Otsu) thresholding methods to segment vasculature. We also tested a random Forest machine learning-based approach and a deep learning approach. The random forest machine learning approach worked with image-derived local features. Intensity, gradient intensity, and local structure were computed at different scales using gaussian blurring. These derived features were then used in random forest classifiers ^14^. The Deep Learning approach utilized the U-Net model specifically designed for biomedical image segmentation ^15^.

The training data for the machine learning-based and deep learning-based vascular segmentation approaches consisted of representative images of segmented vascular labeling from intact tumor light sheet datasets. Roughly 30 two-dimensional (2D) tumor images of segmented blood vessels whose accuracy was evaluated by a biologist were used as ground truth.

Segmentation of blood vessels in the training dataset was performed using ZEN 3.0 software from Zeiss. The representative images used for training were selected across different z planes across the depth of the light sheet datasets to allow machine learning models to see variations in background intensity across different z planes and support accurate segmentation of blood vessels traversing multiple z planes.

To evaluate tumor vascular segmentation results across different approaches, two main testing strategies were utilized. In the first strategy, the vascular segmentation model was trained on two tumor light sheet datasets and tested on a third separate dataset. The second strategy was to use the first half (chronologically for z layers) of all 2D ground truth images from all light sheet datasets for model training and the second half for segmentation accuracy testing. Vascular segmentation masks and ground truth segmentation masks were then compared to quantify model predicted segmentation accuracy using DICE score metrics ^16^.

#### 1.5.2. Tumor boundary segmentation

To characterize drug distribution within the tumor, image segmentation was performed to detect the tumor boundary using data from the Syto16 channel labeling all cell nuclei. This segmentation process generated a 3D mask of the tumor surface. We found that the tumor boundary could be appropriately detected using a thresholding-based segmentation approach. We compared three thresholding methods - Otsu ^17^, Triangle ^18^, and Yen ^19^ . The threshold was calculated individually for all z layers. Next, the median of calculated thresholds across all z-layers was applied to perform segmentation and create a binary mask of the tumor. Further analysis was performed on the pixels within the tumor boundary. Lastly, to support interpretation of drug and vascular tumor distribution, we separated the tumor into two regions of interest, the core, and the periphery, following the work by Dobosz et. al, as an example we assumed that if the tumor were a hypothetical sphere, then the region with a distance to the tumor boundary less than 20% of the radius of the hypothetical sphere could be considered the periphery, and tissue within that could be considered the tumor core ^9^.

#### 1.5.3. Vascular distance map creation

Following blood vessel and tumor tissue segmentation, we computed a distance map to characterize drug penetration from tumor blood vessels into neighboring tissue. In this step each pixel was assigned the distance to the nearest blood vessel. The traditional distance transform operation performed in 2D was not suitable for tumor vasculature since blood vessels typically traversed more than one z plane in our high resolution three-dimensional LSFM datasets.

Therefore, we used the 3D Euclidian distance transform operation. The 3D distance map was calculated utilizing an open-source implementation^20^, which allows for multi thread computing and the package is designed especially for fast processing.

#### 1.5.4. Data aggregation

In the final step of our tumor LSFM data analysis pipeline, the outputs from previous steps were aggregated to support interpretation. Profiles of the drug (oncolytic virus CVA21) penetration away from blood vessels into surrounding tumor tissue were obtained as follows: First, the output from our three-dimensional vascular distance map was discretized. Next, the average fluorescence signal intensities in the CVA21 channel were calculated in every bin, allowing us to plot drug penetration gradients. We compared the drug penetration profiles across the tumor core versus periphery. Blood vessel segmentation results were also used to calculate and compare the fraction of vascular pixels in the tumor core versus the periphery.

## 2. Examining β-amyloid distribution in the brain using tissue clearing and LSFM

### 2.1. Animal model, tissue collection, and processing

All animal experimental protocols were approved by the Institutional Animal Care and Use Committee at Merck & Co., Inc., Rahway, NJ, USA and were carried out in accordance with the NIH Guide for the Care and Use of Laboratory Animals ^11^. TgCRND8 mice (Taconic Inc.) over-expressing mutant human amyloid precursor protein (APP) were used for this study ^10, 21^. At 6.5 months of age TgCRND8 mice were euthanized by transcardial perfusion using the same method described above for the tumor LSFM study. Following perfusion and fixation, the brain was extracted from the skull and split in half along the mid-sagittal plane separating it into two hemispheres. n = 2 hemispheres from separate animals were further processed, imaged, and analyzed to demonstrate our workflow as described below. Tissue clearing was also performed using the same protocol described above for the tumor LSFM study with one key difference – the tissue clearing time from TgCRND8 mouse brain hemispheres in the Smart Clear II Pro device (LCT) was 4 days while using the gentle mode (42° C, 1500 mA). Immunolabeling to detect β-amyloid pathology was performed using a monoclonal rabbit anti human β-Amyloid primary antibody (Cell Signaling Technology, D54D2, Part no. 8243) and Rhodamine Red™-X (RRX) AffiniPure Donkey Anti-Rabbit polyclonal secondary antibody (Jackson Immunoresearch, Part no. 711-295-152) in the Smart Label device from LCT using the same protocol described above for the tumor LSFM study.

### 2.2. Brain LSFM data stitching and visualization

LSFM data was acquired as a mosaic using a 4X objective with an xyz resolution of 1.63 x 1.63 x 4 µm on the UltraMicroscope BLAZE microscope and using the 3.6X objective with an xyz resolution of 1.8 x 1.8 x 4 µm on the Selective Plane Illumination Microscope (SPIM) microscope (SmartSPIM, LCT). Each image z stack for individual xy tile locations was first converted to a single 3D .ims file using the Imaris file converter version 9.7.2 (Bitplane). 3D Imaris files for all xy tile locations were then stitched into a single 3D data file using Imaris stitcher version 9.7.2 (Bitplane). Acquisition meta data stored in the raw OME TIFF files, allowed automatic alignment of 3D z stacks across all xy tile locations in Imaris stitcher. Stitched data was then visualized in Imaris version 9.7.2 (Bitplane).

### 2.3. Brain LSFM data atlas registration

Brain β-amyloid distribution LSFM data was registered to the Allen Brain Atlas ^22^ using an automated process (alignment performed by LCT). In the automated phase, an autofluorescence channel for each brain was registered to 8-20 atlas-aligned reference samples, using successive rigid, affine, and b-spline warping algorithms (SimpleElastix: <u>Source Code</u>^23^). An average alignment to the atlas was created across all intermediate reference brain sample alignments to serve as the final atlas alignment value for the sample.

### 2.4. Brain LSFM data segmentation

β-amyloid segmentation was performed using a custom convolutional neural network created with the Tensorflow python package (LCT, proprietary). The network was based on U-Net and Fully Convolutional Network architectures (<u>Source Code</u> ^24^) to perform a semantic segmentation task. The network outputs a probability map for each 3D input, representing the likelihood that each voxel is part of a beta-amyloid plaque. The probabilities were then thresholded to yield a 3D mask of the same size as the initial image, where positive values indicated detected plaques. Using the previously calculated atlas registration, each plaque location was projected onto the Allen mouse brain atlas to calculate the plaque coverage for each brain region.

## Results

### 1.1. Tissue clearing, labeling, and LSFM enabled *ex vivo* visualization of vasculature and distribution of an OV (CVA21) following intravenous dosing in a xenograft tumor mouse model

**Fig. 4A** shows a static three-dimensional view of a xenograft tumor harvested from an animal 24 hours after receiving a single IV bolus dose of 10^7 TCID50 (Median Tissue Culture Infectious Dose) of CVA21 in saline, processed by aqueous tissue clearing, immunolabeling, and imaged with LSFM. **Supplementary Video 1** plays through the different views and planes across the 3D tumor dataset. The tumor vasculature was immunolabeled using an anti-CD31 antibody and the oncolytic virus was detected using a polyclonal antibody against CVA21. **Fig. 4B & 4C** show two distinct CVA21 distribution profiles at a subcellular level observed via LSFM. In some cells within the tumor environment, CVA21 exhibits a punctate peri-nuclear localization (**Fig. 4B**), while in other instances, CVA21 showed diffused cytoplasmic localization in cells within the same tumor (**Fig. 4C**). Thus, tissue clearing and LSFM not only allowed us to examine the three-dimensional distribution of drug and vasculature across anatomical planes, but also provided spatial information from the macroscopic whole tumor level to the intracellular microscale level.

**Figure 4.**
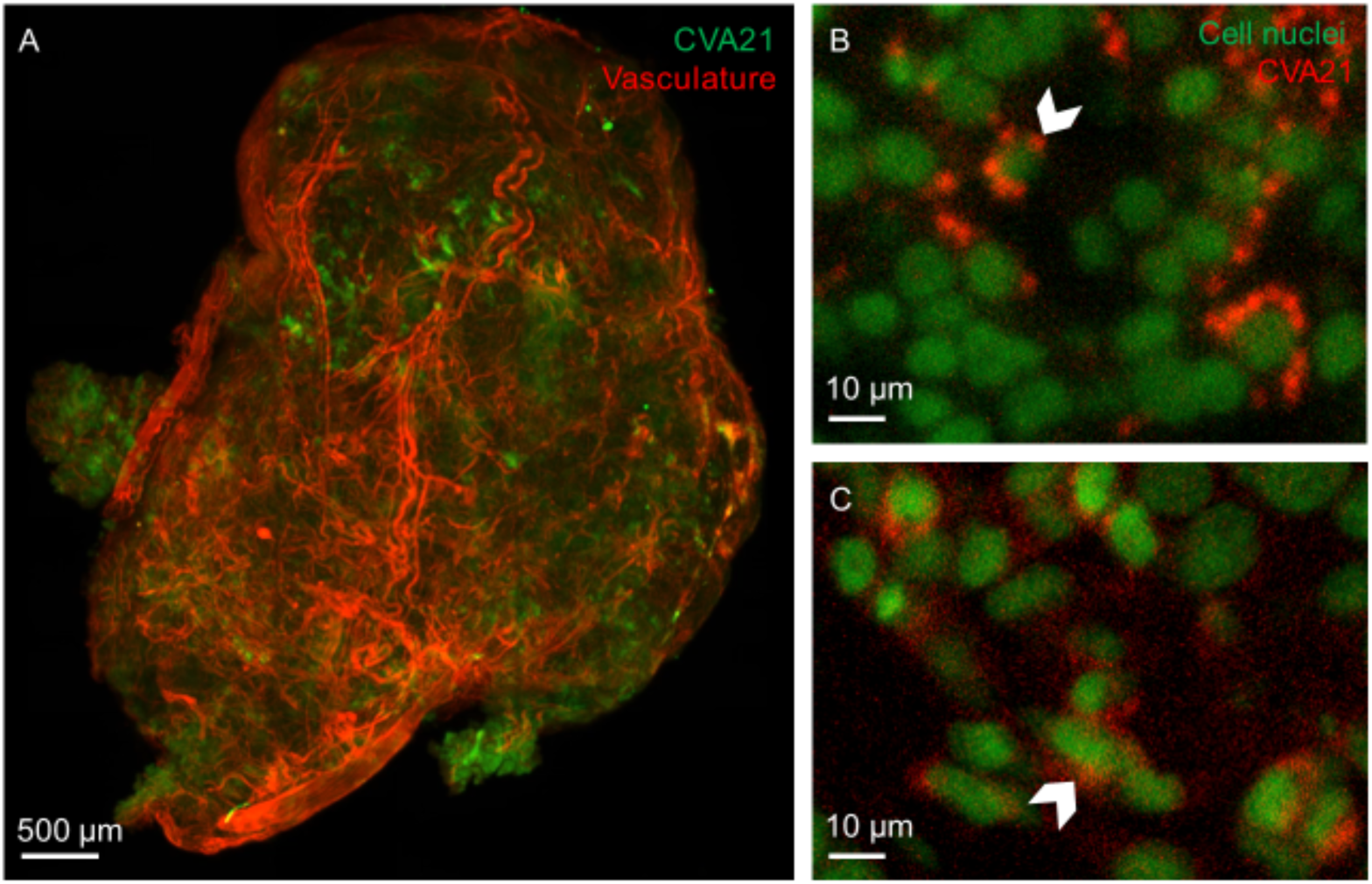
Application of LSFM to investigate oncolytic virus (CVA21) and vascular distribution in xenograft tumors. (A) A whole SK-MEL-28 xenograft tumor 24 hours post intravenous administration of CVA21. Vasculature (red) and CVA21 (green) are detected by immunolabeling. (B) Representative region of interest in tumor LSFM dataset showing punctate subcellular distribution of CVA21 (red) around cell nuclei (green). (C) Representative region of interest in tumor LSFM dataset showing diffuse subcellular distribution of CVA21 (red) around cell nuclei (green).

### 1.2. LSFM enabled estimation of tumor vascular volume

Analysis of LSFM data showed that pixels showing vascular labeling accounted for less than 10

% of the of all pixels within the tumor boundary. Cells within solid tumors often occupy one of two unique microenvironments – the relatively better oxygenated tumor periphery or the hypoxic tumor core ^25–27^. This makes comparisons of drug and vascular distribution across the tumor core and periphery quite informative when trying to understand tumor physiology. For the purpose of our analysis, we assumed the tumor to be a sphere and the outer 20% of the fictional sphere volume to be the periphery ^9^. We observed that the xenograft SK-MEL-28 melanoma tumors across three animals had a nearly two-fold higher volume fraction of CD31 labeled vasculature in the tumor periphery vs the tumor core, although this difference was not statistically significant (**Fig. 5A**).

**Figure 5.**
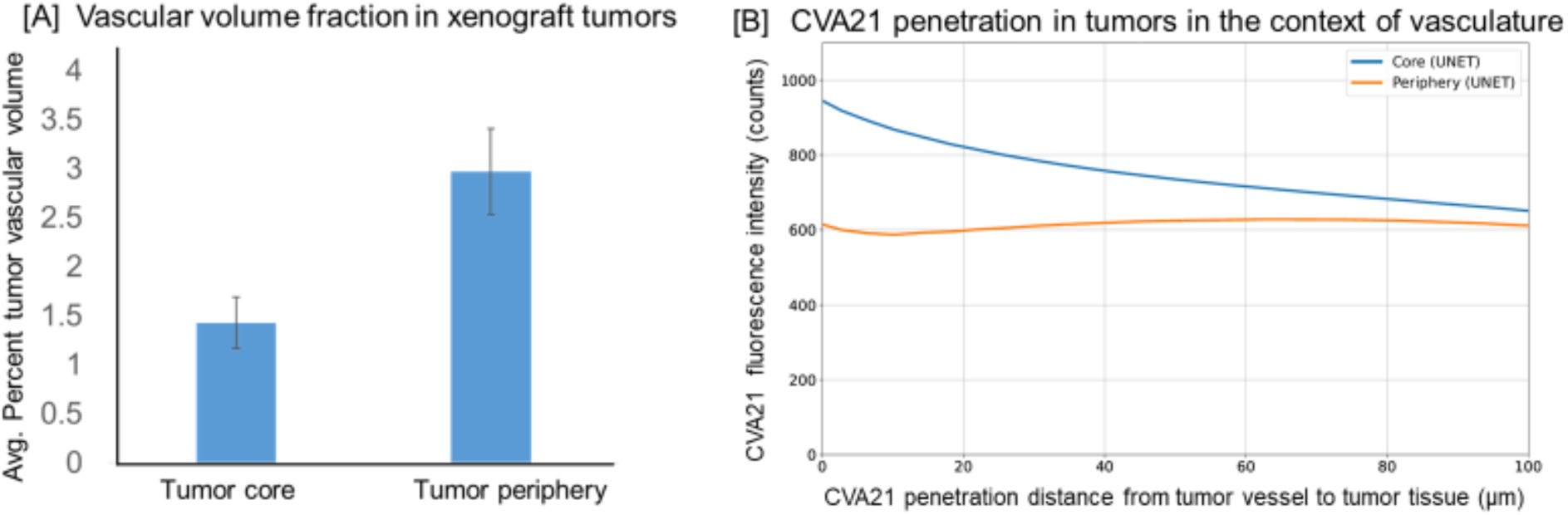
(A) Vascular volume in the tumor periphery and tumor core measured by analyzing LSFM data (n = 3 tumors). The following assumption was used to define the tumor core and periphery: if the tumor were a hypothetical sphere, then the region with a distance to the tumor boundary less than 20% of the radius of the hypothetical sphere could be considered the periphery, and tissue within that could be considered the tumor core (B) CVA21 penetration away from blood vessel walls into tumor tissue.

### 1.3. LSFM enabled evaluation of CVA21 penetration away from blood vessel walls and into tumor tissue

To assess the penetration of intravenously administered CVA21 into tumor, the gradient of viral signal intensity from blood vessel walls up to 100 µm into tumor tissue was measured. The penetration of CVA21 away from blood vessel walls into tumor tissue appeared to be more pronounced in the core versus the periphery (**Fig. 5B**). This suggests that blood vessels in the tumor core are either potentially more permeable due to abnormal vasculature and/or there is enhanced viral replication under hypoxic conditions compared to the periphery ^28^.

### 2.1. LSFM enabled visualization of β-amyloid plaques in intact brain hemispheres of an Alzheimer’s disease mouse model

Preclinical disease models and the ability to assess distribution of pathology are critical for assessing the pharmacology of therapeutic programs. Here, we share an example of using tissue clearing, immunolabeling, and LSFM combined with a unique data visualization approach to understand the spatial distribution β-amyloid pathology in a mouse model of Alzheimer’s disease. TgCRND8 is a transgenic mouse model over-expressing mutant human amyloid precursor protein (APP) ^10, 21^. A static view of the 3D brain β-amyloid plaque distribution dataset obtained via tissue clearing, immunolabeling, and LSFM can be seen in **Fig. 6A**. Representative regions of interest highlight β-amyloid plaque distribution in the cortex (**Fig. 6B**) and hippocampus (**Fig. 6C**). **Supplementary Video 2A** plays through the different views of the 3D β-amyloid plaque distribution light sheet data across the intact TgCRND8 mouse brain hemisphere, while **Supplementary Video 2B** plays through the same TgCRND8 mouse brain hemisphere across serial 2D views of all z planes 4 µm apart. 3D histology allows us to quantify β-amyloid plaque distribution at a resolution comparable to high resolution optical imaging tools such as confocal and two photon microscopies, but now across the intact whole brain hemisphere rather than just a few representative regions.

**Figure 6.**
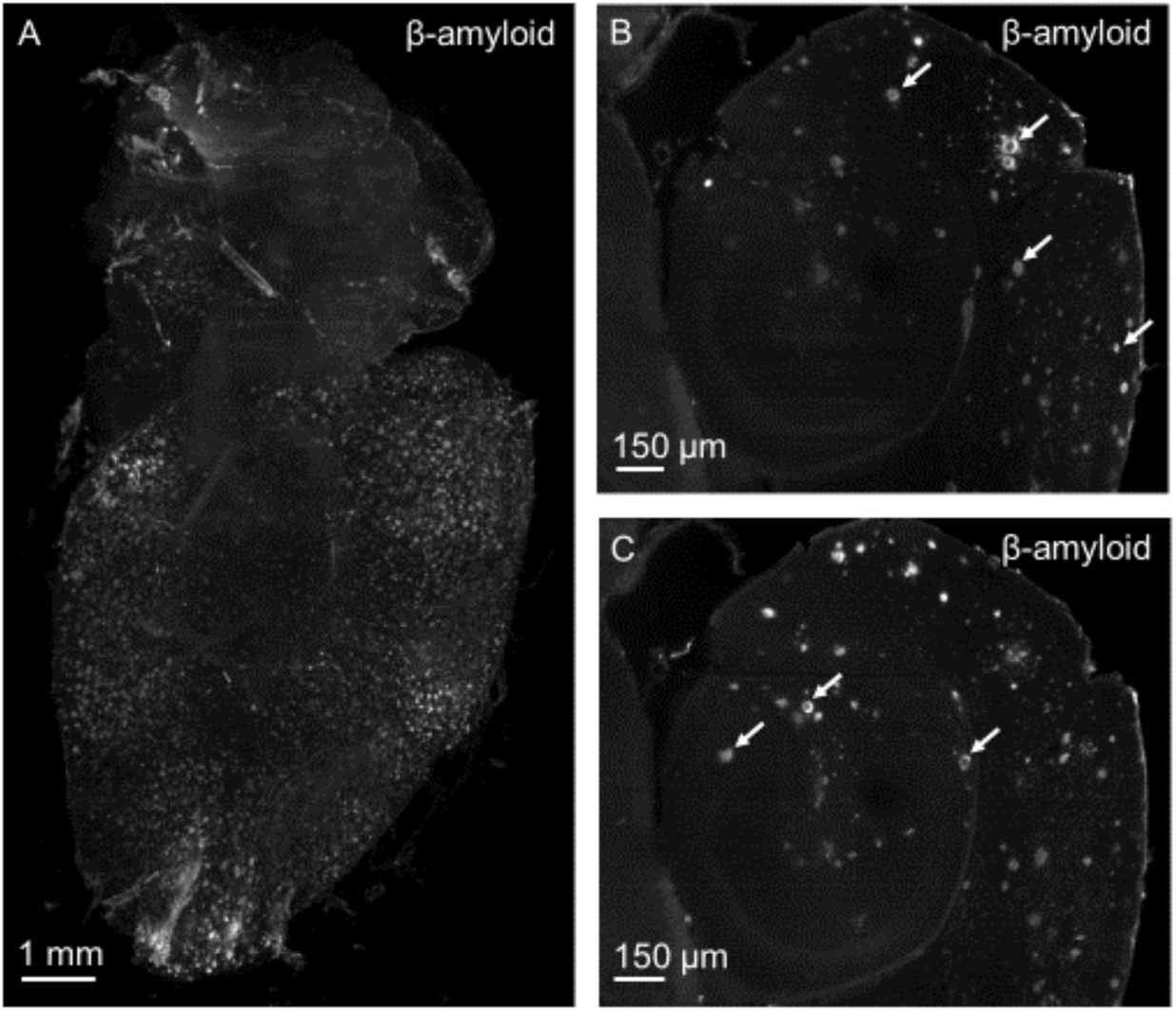
LSFM data demonstrating β-amyloid plaque distribution in the TgCRND8 mouse model of Alzheimer’s disease. (A) Static view of the intact brain hemisphere LSFM data following β-amyloid immunolabeling. (B) Representative region of interest showing β-amyloid plaques in the cortex. (C) Representative region of interest showing β-amyloid plaques in the hippocampus.

### 2.2. β-amyloid plaque density was high in the entorhinal cortex, olfactory areas, and isocortex in the TgCRND8 Alzheimer’s disease mouse model at 6.5 months of age

Following segmentation of LSFM data to detect β-amyloid plaques in the TgCRND8 mouse brain hemisphere dataset, the plaque density was expressed as a heat map registered to the Allen mouse brain atlas using the LCT Smart Analytics platform to demonstrate spatial visualization of plaque data across discrete brain regions (**Supplementary Video 3**). β-amyloid plaque volume in discrete brain regions of interest (ROI) expressed as a fraction of the ROI volume, showed plaque density is high in the olfactory area, isocortex, and entorhinal area in the TgCRND8 AD mouse model at 6.5 months of age (**Fig. 7A**). β-amyloid plaque volume in discrete brain ROIs expressed as a fraction of the total plaque volume in the whole brain hemisphere, showed that the isocortex potentially contributes substantially to the overall β-amyloid plaque burden in the TgCRND8 AD mouse model at 6.5 months of age (**Fig. 7B**).

**Figure 7.**
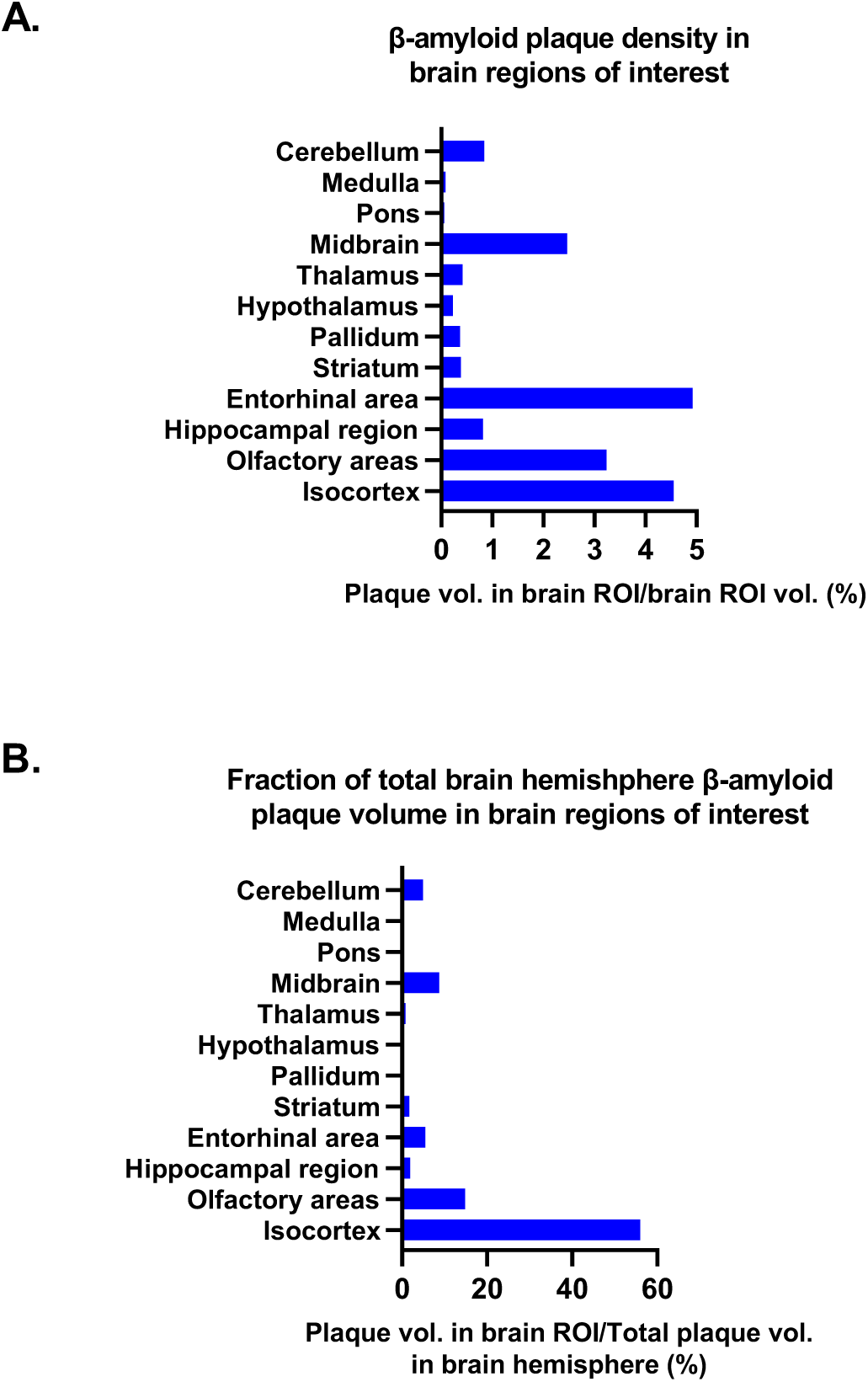
(A) β-amyloid plaque density in the TgCRND8 AD mouse brain regions of interest (ROI) at 6.5 months of age; plaque volume in ROI expressed as a percentage of ROI volume. (B) Fraction of total brain β-amyloid plaque volume in the TgCRND8 AD mouse brain regions of interest (ROI) at 6.5 months of age; plaque volume in ROI expressed as a percentage of total plaque volume in brain hemisphere.

## Discussion

Innovation in tissue clearing and LSFM methods has been instrumental in helping close a key gap in imaging technology – enabling the acquisition of three-dimensional spatial information for whole organs and tumors, without compromising on subcellular resolution and multiplexing. Applications for this burgeoning technology have rapidly spanned spatial assessment of a variety of different organs such as the pancreas ^29, 30^, heart ^31^, kidneys ^32^, brain ^33^, liver ^34^, bone marrow ^35^, and even whole organisms ^36, 37^. Here we explored oncology and neuroscience - two therapeutic areas that could significantly benefit from tissue clearing and LSFM applications to help drive drug discovery. While this work demonstrates workflows and applications of LSFM to investigate the distribution of anatomical and pathological biomarkers relevant to a drug discovery and development setting, the authors acknowledge the potential need for further comparison of LSFM results with traditional sliced tissue immunohistochemistry and optical imaging is warranted.

## 1. Tissue clearing and LSFM will advance how imaging supports oncology drug discovery & development

There has been rapid progress in the development of innovative macromolecular therapeutic modalities and identification of novel targets in oncology over the past decade. However, oncology drug development is impeded by the lack of tools that can achieve 3D scale-spanning spatial interrogation of intact tumors to not only characterize tumor heterogeneity, but also to fully capture the distribution of actionable biomarkers, macromolecular drug, targets, and drug-target engagement dynamics at a cellular resolution. This critical gap can be addressed with tissue clearing and LSFM ^9, 38–40^. In this manuscript, we shared two use cases for LSFM in a drug discovery setting. The first use case described the characterization of tumor vasculature and distribution of the CVA21 oncolytic virus (OV). OVs are viruses that preferentially infect and kill cancer cells and the dying cancer cell releases tumor-associated antigens ^41, 42^. Optimal delivery to the tumor microenvironment (TME) is one of the key challenges facing OV development ^42^. OVs such as the recent FDA approved talimogene laherparepvec (T-VEC) are dosed locally via intratumoral injection (IT), to increase OV concentration in the TME ^42^.

However, IT delivery is clinically cumbersome for dosing deep visceral tumors, and it is not applicable to patients with micrometasases ^42–44^. Although intravenous (IV) administration provides the possibility of targeting tumors inaccessible by IT dosing, its use is complicated by factors that could limit access to the TME, such as rapid dilution in the systemic circulation, host clearance, non-specific uptake by peripheral organs, neutralization by antiviral antibodies or serum proteins, and the failure to extravasate through tumor vessels ^44, 45^. There is currently limited published literature describing the tumor biodistribution of OVs following IT and IV dosing. Bulk sampling methods such as PCR fail to provide spatial information related to OV distribution in tumors and how that relates to tumor vascular heterogeneity. Imaging data providing spatial information for OV distribution at a cellular resolution are scant and obtained using imaging approaches that restrict data to a few representative tumor tissue sections. In this paper, we share how LSFM can be used to examine CVA21 distribution following IV dosing and how that may relate to vascular heterogeneity in intact solid tumors. LSFM datasets provide rich 3D spatial information which could be used to train and accelerate the development of artificial intelligence-based tools for image analysis that could benefit both preclinical research as well as medical imaging ^46^.

## 2. Tissue clearing, and LSFM can help characterize disease progression in neurodegenerative disorders

The anatomical complexity of the brain has put neuroscience at the forefront of LSFM applications ^33^. Understanding neurodegenerative disease progression in the context of brain anatomy and function is critical in the development of therapeutic strategies. Organic tissue clearing methods and LSFM have been successfully applied in previous literature to capture the distribution of β-amyloid plaques in whole mouse brains and archival human brain samples ^47^. In this manuscript we applied aqueous tissue clearing and electrophoretic tools for rapid and uniform tissue clearing and shared an example of quantitative readouts mapped to the Allen brain atlas to support visualization and interpretation of complex 3D data. The trends observed for β-amyloid plaque distribution observed in this study followed trends previously described in the literature ^10^. In the future, LSFM data could be used to characterize the complex interplay between factors that may affect Alzheimer’s disease progression such as microglial plaque barrier formation ^48, 49^ and how they are impacted by therapeutic intervention.

## 3. Machine learning-based tools can help address the LSFM data analysis bottleneck

Successful application of tissue clearing and LSFM in a biopharmaceutical setting requires ongoing exploration of ways to increase throughput and consistency in the workflow. Data acquisition throughput has benefited from commercially available laboratory tools. For instance, the SmartClear and SmartLabel electrophoresis devices (LCT) shorten timelines and increase uniformity during tissue clearing and immunolabeling when using aqueous based methods. The BLAZE light sheet microscope (Miltenyi Biotec.) can accommodate multiple samples at once, enabling batch image acquisition. However, image analysis of LSFM data remains a challenge due to large data size (a single dataset can be ∼ 1 Terabyte or more), data complexity, and diversity in desired quantitative metrics to be extracted. An important result shared in this paper is a tumor LSFM data analysis pipeline that could analyze a ∼ 1 Tb dataset in less than 2 hours when run in a high-performance computing environment. Image segmentation for light sheet datasets can be challenging due to the varying background noise across different z-planes and complexity of three-dimensional structures. Machine learning tools can significantly improve accuracy, reduce bias, and support automation of image segmentation in large imaging datasets ^50, 51^. Using DICE scores, we demonstrated that methods such as random forest and deep learning (U-Net) are capable of providing superior tumor vascular segmentation accuracy compared to intensity-based thresholding (**Table 1**, **Fig. 8**). Moreover, random forest-based machine learning and deep learning-based models allow for a considerable level of flexibility when utilizing manually labeled training data. During the models’ training, we encountered several instances where ground truth data were subjected to a certain level of mislabeling, e.g., under segmentation of manually segmented blood vessels. The ability of pattern learning, and the flexibility of the utilized models allowed for overcoming such inaccuracies within the training data (**Fig. 9**).

**Figure 8.**
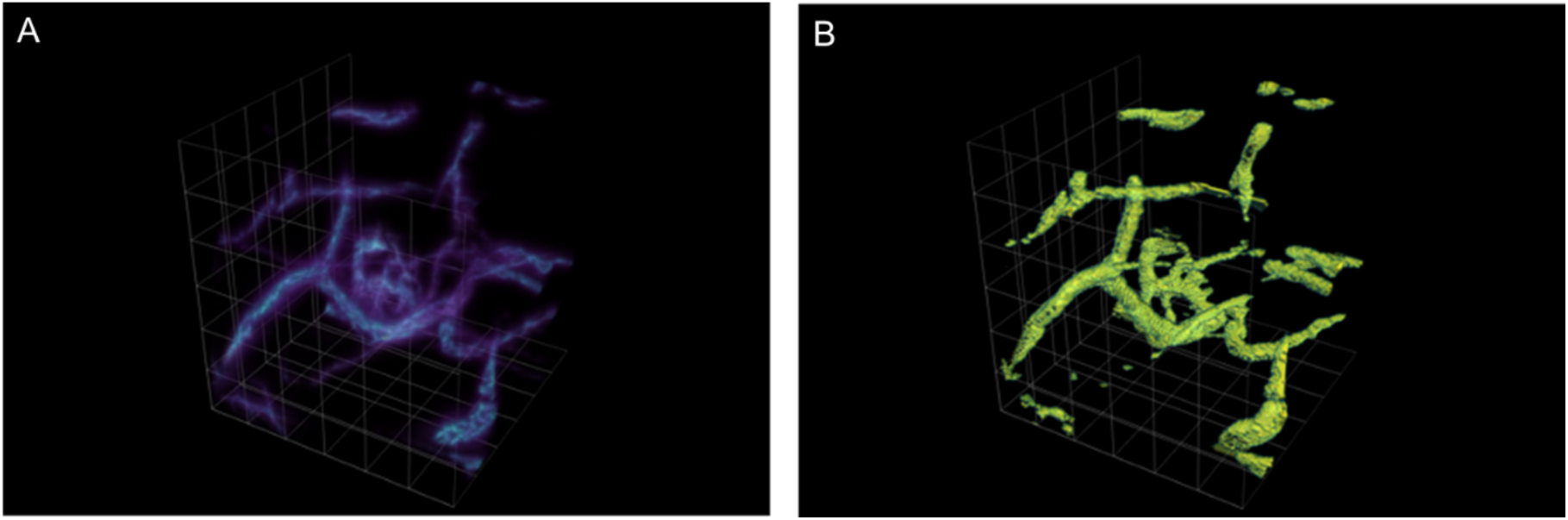
Representative ROI showing (A) tumor vasculature LSFM data and (B) tumor vasculature segmentation results using the U-Net CNN deep learning model.

**Figure 9.**
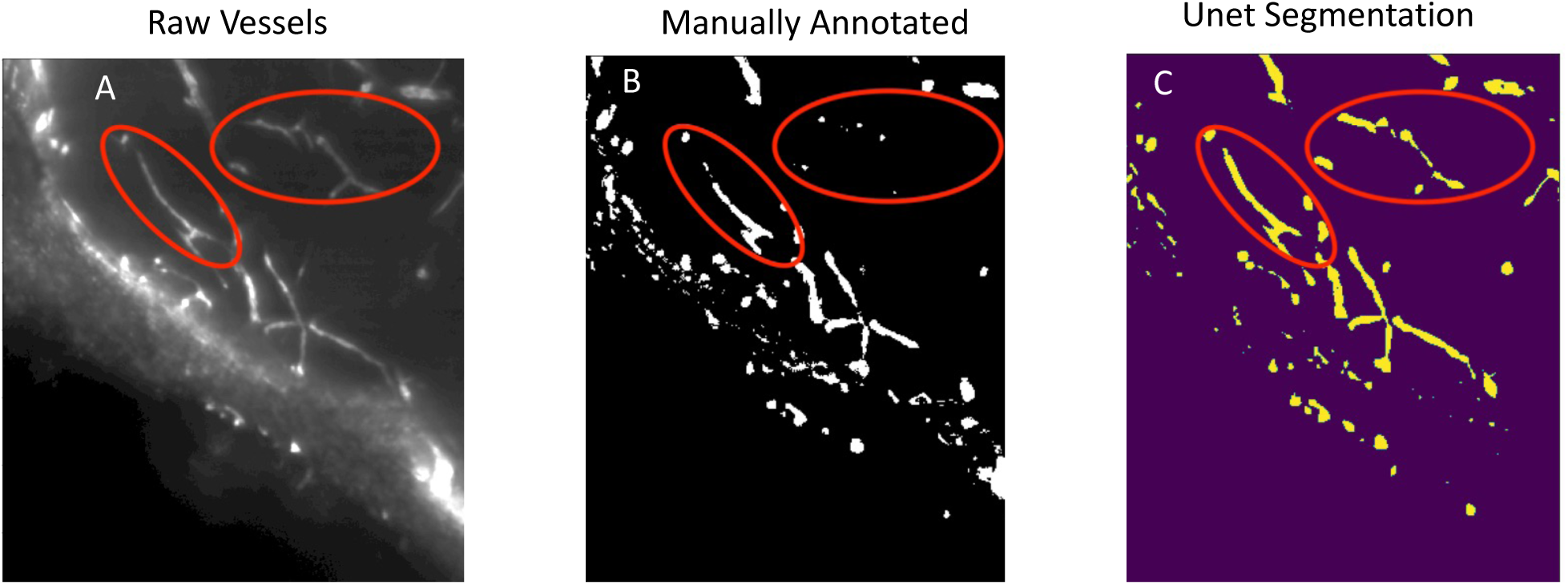
Representative example of vascular segmentation using a machine learning based approach. (A) Raw 2D representative ROI image of vascular labelling from LSFM tumor data. (B) Manually annotated vasculature containing a few under segmented instances (denoted by the red circle). (C) vascular segmentation results obtained by using the U-Net model.

**Figure 10.**
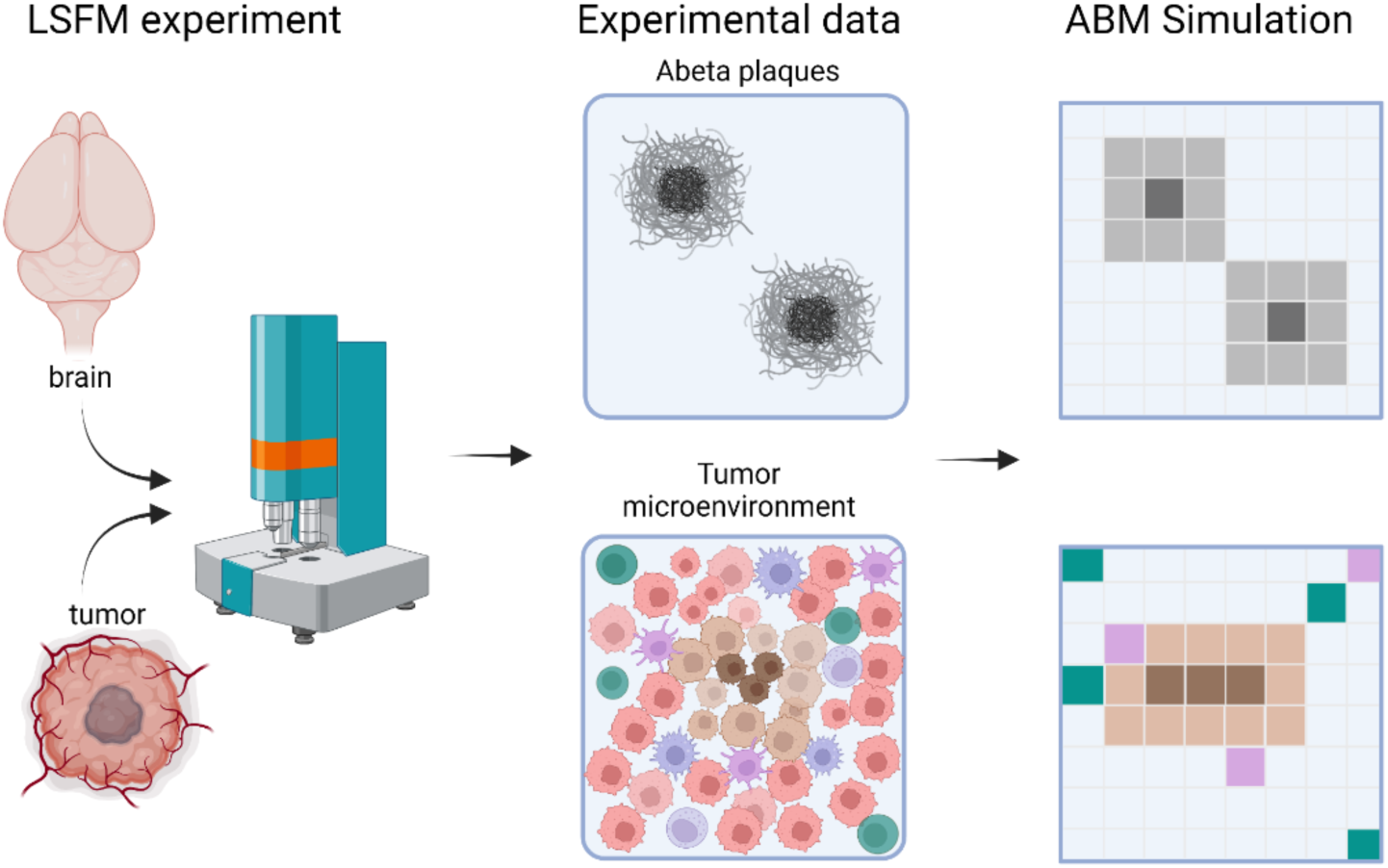
Workflow depicting the combination of LSFM data with agent-based modeling. The spatiotemporal progression of disease pathology can be experimentally obtained with LSFM and described mathematically using an agent-based model (ABM). We have depicted brain and tumor LSFM experiments and the generation of data on Aβ plaques and cells in the tumor microenvironment respectively. Aβ aggregation from monomers to plaques is represented from shades of light gray to dark gray. Tumor microenvironment consists of cancerous cells in pink, necrotic areas in brown, and immune cells in purple and blue. Illustrative examples of ABM simulations that could describe these disease processes are shown on the right. Created with BioRender.com.

**Table 1.**

| Training datasets<br>(sample IDs) | Testing datasets<br>(sample IDs) | Random<br>Forest | U-Net | Thresholding |
| --- | --- | --- | --- | --- |
| 7, 8 | 5 | 0.72 | 0.74 | 0.62 |
| 5, 8 | 7 | 0.8 | 0.76 |  |
| 5, 7 | 8 | 0.79 | 0.77 |  |
| ½ of the images<br>from all light sheet<br>datasets | Second half of<br>images not used<br>for training | 0.79 | 0.79 |  |

## 4. LSFM data could support model informed drug discovery and development

LSFM data can be used to support quantitative systems pharmacology (QSP) and physiology-based pharmacokinetics (PBPK) modeling to inform drug development. For example, a mechanistic in-silico predictive model has been used to simulate CVA21 viral kinetic and dynamic processes following IT and IV delivery ^52^. In this model, the tumor compartment was divided into vascular and tissue interstitium sub-compartments. However, the model did not account for differences between the tumor core and tumor periphery, which can be architecturally and physiologically different in solid tumors^26, 27, 53, 54^. Experimental data from LSFM could be used to inform model parameter estimates needed to guide such characterization, and potentially other biological process, such as immune cell infiltration, in both preclinical tumor models and patient derived biopsies^40^. Moreover, such information could support drug design and dose selection, as well as strategies for development of monotherapy and combination therapy strategies.

Drug development in neuroscience is challenging, which is exemplified by the low probability of success for drug approval relative to other fields. QSP models could help to fill this translational gap and support the development of effective disease-modifying therapeutics for many unmet medical needs, such as Alzheimer’s disease (AD).^55^ However, there are often simplifying assumptions introduced into these models due to the lack of data, which limits their applicability and interpretability. Agent-based modeling (ABM) has emerged as a powerful mathematical modeling technique to describe a wide range of natural and biological systems ^56–58^. Simply, ABMs consist of agents that can traverse a region of space over a period of time according to a set of defined rules. In comparison to ordinary differential equation-based models, ABMs are spatially resolved and can account for cellular heterogeneity ^59^. An ABM could be developed and paired alongside an LSFM workflow to provide enhanced spatiotemporal granularity in disease progression models. For example, an ABM LSFM workflow could be applied to describe the progression of amyloid pathology in the brain and cellular heterogeneity in tumors. The ABMs could for instance provide quantitative predictive insights into biological processes governing amyloid dynamics and the role of microglia in plaque formation in the AD brain or the migration of immune cells in tumors. Ultimately, these hybrid experimental-computational workflows can be utilized for the optimization of new drugs. The ABM could provide quantitative insights into biological processes governing amyloid dynamics, the role of microglia in plaque formation, and evaluating the pharmacodynamic effects of therapeutic interventions ^60^.

## Conclusion

In conclusion, this work outlines two examples of the application of tissue clearing and LSFM to evaluate macromolecular drug and biomarker distribution in intact preclinical brain and tumor samples. This study describes the utility of LSFM in drug discovery, novel machine-learning based approaches for LSFM data analysis, and discusses the broader applications for how such data may be used to support predictive model informed drug discovery.

## Acknowledgements

Cora Ames, Dominic Mangiardi, and Aparna Nambiar supported services provided by Life Canvas Technologies. Renee Gentzel supported technology testing and onboarding at Merck & Co., Inc., West Point, PA, USA.

## Author contributions statement

N.K. conceived the experiment(s), conducted experiment(s), performed data analysis, and wrote the manuscript. P.H., M.V., J.S., and R.P., conceived the data analysis strategy, performed data analysis, contributed towards writing the manuscript, and reviewed the manuscript. N.P. conducted experiment(s), contributed towards writing the manuscript, and reviewed the manuscript. R.M. conducted experiment(s) and reviewed the manuscript. T.F. and P.B. contributed towards writing the manuscript and reviewed the manuscript. S.B. reviewed the manuscript. C.P., C.C., and M.C. reviewed the manuscript and provided mentorship.

## Data Availability statement

Tumor data analysis pipeline code can be found in our GitHub repository (https://github.com/Merck/3D_Tumor_Lightsheet_Analysis_Pipeline). A subset of the raw LSFM datasets are publicly available with an interactive demo image analysis notebook.

(https://github.com/Merck/3D_Tumor_Lightsheet_Analysis_Pipeline/blob/main/demo_script.ipynb).

## Funding information

Research was funded by Merck Sharp & Dohme LLC, a subsidiary of Merck & Co., Inc., Rahway, NJ, USA.

**Supplementary video 1.** 3D visualization of an SK-MEL-28 xenograft tumor collected 24 post intravenous administration of CVA21 and subjected to tissue clearing, immunolabeling, and LSFM.

**Supplementary video 2.** (A) 3D visualization of β-amyloid plaque distribution in a brain hemisphere of a 6.5-month-old TgCRND8 mouse following tissue clearing, immunolabeling, and LSFM. (B) 2D visualization across all z planes at a step size of 4 µm of β-amyloid plaque distribution in a brain hemisphere of a 6.5-month-old TgCRND8 mouse following tissue clearing, immunolabeling, and LSFM.

**Supplementary video 3.** β-amyloid plaque density in the TgCRND8 mouse brain hemisphere obtained from LSFM data expressed as a heat map registered to the Allen mouse brain atlas using the Life Canvas Smart Analytics platform to enable spatial visualization of quantitative data across discrete brain regions.

